# DisGUVery: a versatile open-source software for high-throughput image analysis of Giant Unilamellar Vesicles

**DOI:** 10.1101/2022.01.25.477663

**Authors:** Lennard van Buren, Gijsje Hendrika Koenderink, Cristina Martinez-Torres

## Abstract

Giant Unilamellar Vesicles (GUVs) are cell-sized aqueous compartments enclosed by a phospholipid bilayer. Due to their cell-mimicking properties, GUVs have become a widespread experimental tool in synthetic biology to study membrane properties and cellular processes. In stark contrast to the experimental progress, quantitative analysis of GUV microscopy images has received much less attention. Currently, most analysis is performed either manually or with custom-made scripts, which makes analysis time-consuming and results difficult to compare across studies. To make quantitative GUV analysis accessible and fast, we present DisGUVery, an open-source, versatile software that encapsulates multiple algorithms for automated detection and analysis of GUVs in microscopy images. With a performance analysis, we demonstrate that DisGUVery’s three vesicle detection modules successfully identify GUVs in images obtained with a wide range of imaging sources, in various typical GUV experiments. Multiple pre-defined analysis modules allow the user to extract properties such as membrane fluorescence, vesicle shape and internal fluorescence from large populations. A new membrane segmentation algorithm facilitates spatial fluorescence analysis of non-spherical vesicles. Altogether, DisGUVery provides an accessible tool to enable high-throughput automated analysis of GUVs, and thereby to promote quantitative data analysis in GUV research.

## Introduction

Giant unilamellar vesicles (GUVs) are aqueous compartments enclosed by a lipid bilayer membrane. ^1^ Since their diameter is typically between 5 and 100 micrometers, which is comparable to the size of eukaryotic cells, and their membrane is composed of phospholipids just like plasma membranes, GUVs are considered a good model system for living cells. As such, GUVs have gained great interest from researchers in biochemistry, biophysics, synthetic biology and applied medicine.

One of the most classical applications of GUVs is in studying the physicochemical properties of biological membranes. Being larger than other biomimetic membrane systems such as large unilamellar vesicles (LUVs) and small unilamellar vesicles (SUVs), GUVs can be easily observed with optical microscopy. Accordingly, GUVs are extensively used to study a wide variety of membrane properties: mechanics, ^2^ lipid diffusivity,^3^ permeability,^4^ as well as lipid order ^5^ and domain formation. ^6, 7^ Moreover, GUVs are often used to study the biophysical mechanisms that underlie important cellular events such as membrane growth,^8^ budding, ^9^ fission^10^ and fusion. ^11^ Furthermore, GUVs provide a suitable chassis in the endeavour of building synthetic or artificial cells. ^12–14^ In this emerging research field, cellular functionalities are being reconstituted from chemical or biological building blocks with increasing complexity, with the eventual goal to understand the minimal requirements for life at the cellular level. ^15^ Lastly, the biocompatibility of GUVs makes them also interesting in the context of targeted drug delivery.^16^ They overcome size limitations of the SUVs that are typically used, offering a way to deliver more cargo per particle.

As GUVs are becoming a widely used tool in synthetic biology, also the possibilities for their production are growing. By now, numerous production methods have been developed to produce GUVs, ranging from simple and quick bulk methods with low-cost equipment to advanced microfluidic methods. Two major pathways can be distinguished for the production. First, GUVs can be formed by hydration of a dried lipid film, either by spontaneous swelling on solid supports or porous substrates, or by application of an electric field. ^1, 17–19^ Second, GUVs can be templated from water-in-oil emulsion droplets, for example by the inverse emulsion method, ^20, 21^ with microfluidics^22, 23^ and by continuous Droplet Interface Crossing Encapsulation (cDICE).^24, 25^ With the versatile options for formation, the design possibilities for GUVs have become legion: from simple membranes composed of a single lipid type to complex biological lipid extracts, ^26, 27^ charged^28, 29^ or bio-functionalized membranes, ^30^ membranes with asymmetric leaflets^31^ or including membrane proteins, ^32^ in physiological buffers,^29, 33^ encapsulating functional proteins^8, 25^ or even active matter.^34, 35^

By far the most widely used characterization technique for GUVs is optical microscopy. GUVs can be imaged in bright field or by fluorescence microscopy upon inclusion of dyes, either membrane-bound or encapsulated. While most studies with GUVs involve simple wide field or confocal microscopy, also superresolution microscopy^36^ and bulk analysis with multi-well plate assays^21^ and fluorescence-activated single cell sorting (FACS)^37^ have been employed. For all the GUV applications described above, it is crucial to evaluate the quality of the produced GUV samples because the success of GUV formation and the resulting vesicle properties can vary substantially dependent on experimental conditions. GUV analysis comes itself with the challenge that in most reconstitution experiments, the formed vesicles are polydisperse in size, shape, the presence of membrane structures and encapsulated content. The complex appearance of heterogeneous GUV populations therefore demands a quantitative characterization by accurate descriptors and robust statistics.

Despite the experimental ease of producing and imaging GUVs, their quantitative image analysis has received relatively little attention. ^38^ Typically, GUVs are either manually detected in the image and afterwards (manually) processed to extract data,^30, 39–44^ or custom-made scripts are used to process specific data sets and generate a pre-defined set of output parameters.^21, 25, 45–47^ Consequently, GUV image analysis is currently time-consuming and non-standardized, making it difficult to directly compare the outcome of different studies.

In the field of cell biology, analysis workflows do exist for automated characterization of cell or tissue image data, combining standardized detection modules with reporting a multitude of output variables. ^48, 49^ Unfortunately, these analysis workflows offer a limited compatibility with GUV data sets. While cells generally have a complex morphology, GUVs are typically near-spherical, highly symmetric 3D objects. Due to their often predictable shapes and intensity profiles, rapid and efficient detection and characterization of vesicles benefit from a simplified approach. Furthermore, irrespective of the application in which GUVs are used, the same set of descriptors are typically of interest, in particular vesicle size, shape, membrane intensity (lamellarity), and spatial intensity profiles of GUV membrane and content.

Some examples of openly available GUV analysis software do exist, laying the ground to make large-scale GUV analysis more accessible. However, they are all either geared towards specific, predefined analysis (membrane permeability,^50^ heterogeneity in membrane signal ^51^) or they have limited compatibility with input data sets and vesicle types (confocal microscopy images, ^46^ spherical vesicles ^52^), requiring high signal-to-noise ratio of the membrane or predefined vesicle shapes for vesicle detection. In addition, most available software lacks a user-friendly interface that allows for interactivity during the detection and analysis procedures, in turn imposing a steep learning curve on new users.

To meet the requirement for accessible and flexible quantitative vesicle analysis, we have developed DisGUVery, an open-source software for the analysis of GUVs in microscopy images. Our tool encapsulates multiple algorithms for the detection of vesicles and the subsequent analysis of their morphology and content under a Graphical User Interface (GUI) based on Python. The software is designed to allow for maximal flexibility in data input, processing and analysis, enabling the user to work with a variety of imaging sources, to export variables of interest at any point during the processing, and to choose between a set of pre-defined detection and analysis modules. Our toolbox provides a general, fast, and user-friendly approach towards quantitative and high-throughput GUV sample characterization, which should be of broad use for the fields of membrane biophysics, cell biology, and bottomup synthetic biology.

## Results and discussion

The general workflow of DisGUVery is summarized in fig. 1A. The starting point is a microscopy image with GUVs visible either via fluorescent labelling (epifluorescence or confocal microscopy as shown in fig. 1B) or in bright field microscopy (phase contrast images). In case of multi-channel images, it is possible to select the channel that should be used for GUV detection, typically, but not limited to, a channel where a membrane dye is imaged, or a phase contrast channel in which vesicles are clearly visible. To allow for high-throughput analysis of large GUV data sets, images can be processed in batches both for vesicle detection and subsequent analysis. When processing is done, detection results can be inspected in the software, and erroneously detected vesicles or unsuccessfully processed images can be discarded before further analysis.

**Figure 1:**
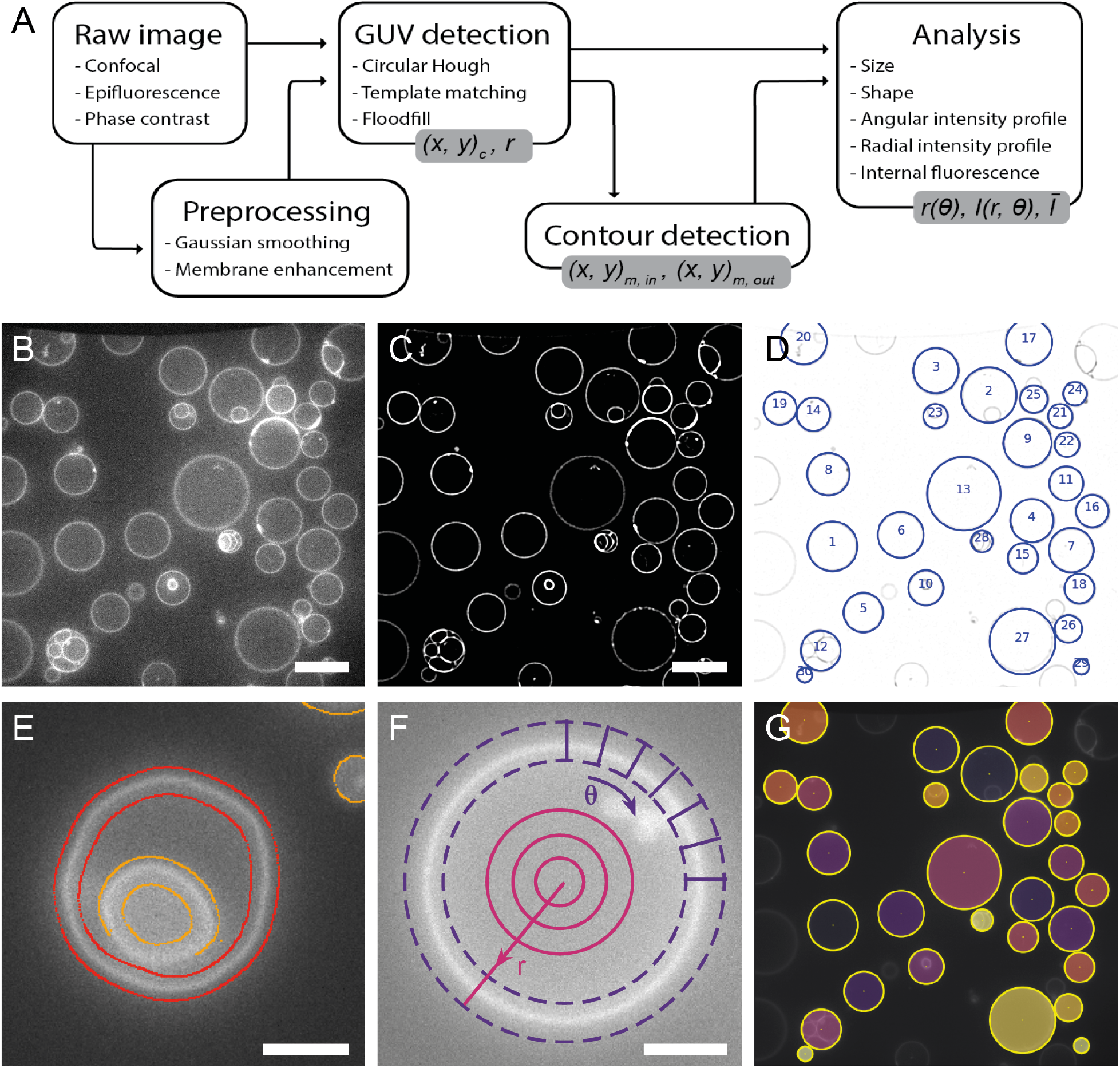
General workflow of GUV detection and analysis by DisGUVery. (A) Visual representation of the workflow. Output variables are shown in grey boxes. (B) Example of an unprocessed single-plane confocal fluorescence microscopy image of GUVs, used as input for the analysis. Scale bar is 20*µm*. (C) Processed image after enhancement of the membrane signal. (D) Vesicles detected by Circular Hough Detection indicated with blue circles and object index number. Contrast is inverted for visualization. (E) Refined contour detection distinguishes the enclosing membrane of a detected vesicle (red) from other fluorescent structures in the image (orange). Scale bar is 5*µm* (F) Radial (magenta) or angular (purple) intensity profiles can be extracted from detected GUVs. Scale bar is 5*µm* (G) Masks (colored) can be created from detected vesicles (yellow circles) to extract internal fluorescence of vesicles.

Prior to vesicle detection, background noise can be reduced by processing the input image using a Gaussian smoothing filter followed by an enhancement of the membrane signal (fig. 1C), both of which can be tuned according to the input image. After this optional preprocessing of the image, GUV detection can be done using one of three different methods, all of which yield the indexed locations and sizes of detected GUVs (fig. 1D). To optimize detection, input parameters can be varied in the GUI and the results can directly be inspected, and wrongly assigned vesicles can be manually discarded. Size distributions can at this point be directly computed and visualized, or other, more complex analysis can be pursued.

For many analysis purposes, such as obtaining vesicle shape descriptors of deformed GUVs^40^ or probing membrane colocalization of fluorescent proteins, ^53^ the precise membrane location is required. We have implemented a membrane segmentation algorithm that can track the membrane of non-spherical vesicles, detecting both the inner and outer edge of the membrane (fig. 1E). When high spatial accuracy is not required, users can also make use of the computationally cheaper basic membrane analysis feature, where the contour from vesicle detection is simply expanded with a certain width to create a ring that contains the membrane (fig. 1F, purple dashed lines). Regardless of the chosen method for membrane segmentation, the angular intensity profiles and the radially integrated intensity profiles can be extracted (fig. 1F), for example to retrieve the angular profile of membrane fluorescence,^54, 55^ the angular profile of a fluorescent membrane-binding molecule, ^56^ or to quantify membrane localization of an encapsulated molecule ^30^ or an externally added membrane-binding protein.^8^

Besides the radial and angular intensity profiles, which provide information about the spatial distribution of fluorescent probes, the average intensity of the vesicle lumen can also be extracted for detected vesicles using the Encapsulation Analysis module (fig. 1G). Analysis of internal fluorescence is essential for studying the efficiency of encapsulation of molecules and other components,^25, 57^ for permeabilization assays where transport of a fluorescent probe across the membrane is tested,^58–60^ and for fluorescence-based measurements of the activity of internal metabolic pathways.^8, 61^

The analyses mentioned above are some of the methods that we have predefined in the software. However, we want to stress that the use of DisGUVery is not limited to these analyses. Since it is possible to export results from object detection, contour detection, and from the analyses at any point in the process, users can extract the relevant information and perform their own analyses outside the software.

### Vesicle detection

We have implemented three different methods for the detection of vesicles in microscopy images: Circular Hough Transform (CHT), Multiscale Template Matching (MTM) and Floodfill detection (FF). As the underlying principle for object detection is different for each of the methods, they allow detection of a variety of vesicle shapes and imaging sources. The first method, based on the circular Hough transform of the object edges, ^62^ is commonly used in the detection of GUVs as it recognizes circular objects with little influence of the intensity profile.^50, 52^ As a result, detection by CHT depends mostly on vesicle shape and not on image intensity, providing a robust method with a high selectivity towards circular vesicles. When the vesicle shape is not circular, but is predictable, for example, by having a population of similar looking vesicles in an image, detection can be done via the second method: template matching. ^63^ We have implemented a slight variation of this method, Multiscale Template Matching (MTM), by allowing the re-scaling of the template to multiple sizes. MTM works by the convolution of the image with a target object, or template, which can be an image of a typical vesicle. Regions in the image are then assigned as detected objects when this template matches the region, with the scaling of the template enabling the size-invariant detection of vesicles. The third method, Floodfill detection (FF), is based on an absolute intensity difference between membrane and background signals. ^64^ By thresholding the image, membranes can be distinguished from the background and closed membrane contours in the thresholded image are assigned as vesicles. Floodfill detects vesicles based on membrane fluorescence, regardless of their shape. Note that FF has been implemented previously for vesicle detection by Blanken et al., ^46^ but with a different starting point for the floodfill algorithm (the seed point). While their algorithm floods all the regions within GUVs by scanning a range of intensity thresholds and seed points, ours floods the surrounding back-ground, which has the computational advantage of using only a single thresholding intensity and a single seed point.

We evaluated the performance of the three vesicle detectors on different types of microscopy images: fluorescence confocal, epifluorescence, and phase contrast. We focused on two main aspects to determine the quality of the detectors: how good are they at detecting vesicles within an image? And, how sensitive is this detection to different factors, e.g., detector parameters or image source? Detection outcomes of the software were benchmarked against human visual detection. We started by optimising the detector parameters on a single image. Figure 2A-C shows an example of the detection results for all detectors on a single confocal image. In this case, the optimal parameters are those which allow the detection of the highest number of vesicles in the image, regardless of their characteristics, while avoiding artefacts in the detection. Once this optimisation has been done, we explore the parameter space of each of the detectors and evaluate the F_1_-score, ^65^ defined as:

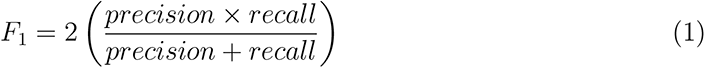

**Figure 2:**
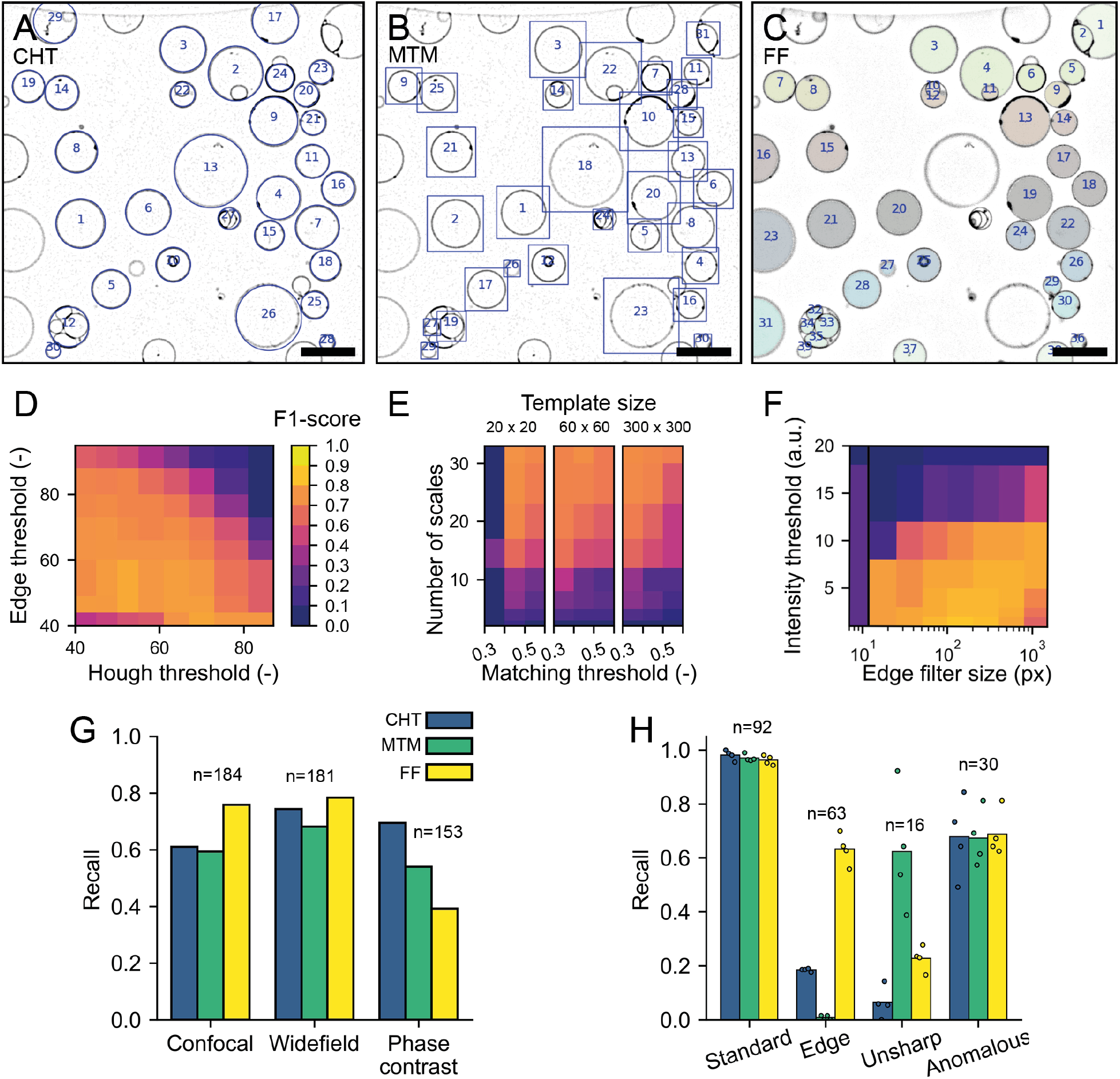
Performance of Vesicle Detection. (A-C) Detected vesicles with the Circular Hough Transform (A, blue circles), Multiscale Template Matching (B, blue bounding boxes) and Floodfill (C, colored objects). The contrast of the images has been inverted for visualisation purposes. Scale is 20*µm* in all images. (D-F) F_1_-score of CHT (D), MTM (E) and FF (F) for different parameter values (see methods for details on the parameters). The color scale in (E-F) is the same as in (D). (G) Recall of vesicle detection for confocal fluorescence, widefield fluorescence and phase contrast images using the three different detectors. (H) Detection recall for different subcategories of vesicles as performed by four different human observers. See main text for more detailed explanation of the use of categories. Individual data points represent results of the individual observers, bars represent average recall values.

Here, 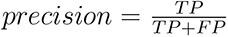 and 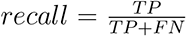 with *T P*, *FP* and *FN* being true positives, false positives and false negatives, respectively. In this study, true positives are detection results that correspond to vesicles, false positives are identified objects that are not vesicles, and false negatives represent GUVs that have not been detected. The reference human visual analysis was performed by a single observer by counting all GUVs in the images, regardless of vesicle size, type, appearance, or position in the image. GUVs at the edge of the image were included as long as a part of the membrane was visible.

We have chosen to use the F_1_ score as an output metric to evaluate our detectors, because it is mainly penalized by false negatives and false positives, both of which are useful output parameters in object detection. As such, F_1_ is amply used in object detection problems. ^66^ Since the F_1_ score is the harmonic mean of precision and recall, both are weighed equally into a single output. While recall represents the fraction of objects in the image that are detected, precision denotes which fraction of detected objects are vesicles. Figure 2D-F show the F_1_ scores as a function of pairs of critical parameters inherent to each of the detectors, with the exception of Floodfill (FF). For FF detection, we chose instead the size of the filter for membrane signal enhancing in the pre-processing step, as we have found it to be critical for the method performance (fig. 2F). We find that all detectors show a region within their parameter space in which the F_1_ score is maximum and their performance is best. Note that the F_1_ score only changes within 10% of its maximum value for a large set of parameters, suggesting that a precise optimisation of the parameters is not necessary, which facilitates batch-processing of data sets with similar images. Notably, CHT appears to be the least sensitive method to changes in the parameter choice of threshold values (fig. 2D), with performance dropping only at large thresholds (*∼* 50% increase from the optimal value). For

MTM, the number of scales used to resize the template is the critical factor in achieving a good performance. This is strongly dependent on the dataset used, more specifically on the polydispersity in vesicle sizes. For the wide distribution of vesicle sizes in our sample, a large number of template scales allows precise matching of the template with GUVs of different sizes. To ensure that the size of the template prior to scaling does not play a crucial role in detection, for example due to pixelation effects, we tested templates of different sizes which resulted in similar performance (fig. 2E). As expected, FF detection depends greatly on the intensity threshold (fig. 2F), with higher values not allowing the vesicles to be properly segmented in the binary mask of the thresholded image. Interestingly, in the images tested, the pre-processing step of membrane enhancement is crucial for FF detection to succeed, as without it, FF detection simply fails to detect vesicles as is demonstrated in fig. 2F at the smallest edge filter size.

We then investigated the extent to which the imaging conditions impact the detection methods, for example, by changing the type of microscopy used to visualize the vesicles. We compiled a dataset of 5 images for each one of the three standard microscopy techniques mentioned above, resulting in a total of over 200 vesicles for each imaging method. For each dataset, we perform the detection using the parameters that were finely tuned for a random image within the set. To measure the performance of the detectors, we evaluate separately the precision and the recall. The reason for this split being that all detectors show a high precision value (between 0.9 and 0.99) for the different data sets (fig. S1), with any differences in the detection performance being represented predominantly in the recall metric. In fig. 2G it can be seen that in both confocal fluorescence and epifluorescence images, all detectors are able to detect vesicles with a recall between 0.6 and 0.8, meaning that 60 to 80% of vesicles are properly detected. However, for phase contrast images, recall decreases, and significantly so for Floodfill detection (to 40%). The low performance of the FF detector in phase contrast is expected based on the intensity profile that vesicles show in this type of microscopy, where the GUV membrane does not represent an intensity maximum but instead the steepest intensity gradient. Furthermore, intensity variations either due to inhomogeneity of illumination or to the presence of surrounding objects interferes with detection based on an absolute intensity threshold. Unsurprisingly, CHT is the most robust method across imaging sources for the sample we investigated, emphasizing its dependence on vesicle shape over intensity profile.

Given that the detectors used target different types of objects, we next looked into what kind of vesicles were being detected by each method. We therefore manually divided all vesicles analyzed in fig. 2G into four categories: vesicles that were located at the edge of the image (‘edge’), vesicles that were out of focus (‘unsharp’), anomalous vesicles (‘anomalous’) and vesicles that do not belong to any of the first three categories which we called ‘standard’ vesicles. A gallery of example vesicles for all subcategories can be found in fig. S2. Anomalous vesicles could be vesicles with a bright membrane signal in their lumen, with very heterogeneous membrane signal, or vesicles that were stuck together in aggregates. Typically, the standard vesicles are those needed for further analysis. In fig. 2H, we can clearly see that while ‘standard’ vesicles are detected similarly by all methods, with recall being nearly 1.0, limiting cases such as vesicles on the edge of the image and those that are out of focus can be easily detected or filtered out depending on the method of choice. CHT and MTM filter out most vesicles at the edge. Furthermore, CHT also misses the unsharp vesicles, which are not detected by FF either. Interestingly, all detectors perform similarly for the anomalous vesicles, with more than 60% of them being detected. Together, these results show the robustness of the different detectors, allowing detection to be performed on a variety of vesicle types and imaging sources.

To illustrate the range of applications of the three detectors for synthetic cell research, we performed vesicle detection in a selection of proof-of-principle experiments. First, we investigated if our methods could be used to track GUV size and number during swelling-based GUV formation. GUV swelling methods include electroformation and gel-assisted swelling and are by far the most popular formation methods, as they are fast and easy and yield large numbers of GUVs.^19, 38, 67, 68^ In these experiments, lipids are first dried on a surface, which can be a hydrogel, an electrode, glass, teflon or a porous material. Subsequent addition of a swelling solution leads to swelling of the lipid film and formation of large numbers of GUVs that are closely packed above the swelling surface (fig. 3A, top image). Using CHT detection, we efficiently detect spherical vesicles even at high surface coverage (fig. 3A, bottom image). By automated detection of GUVs during their formation, growth kinetics could easily be obtained. Since detection by CHT relies on vesicle shape rather than intensity, the method is largely insensitive to touching vesicles or high background fluorescence, both of which are more likely at high packing density. Furthermore, the ability to specify a minimum and maximum GUV radius prevents detection of false positives in dense samples. In addition to GUV production by swelling, detection at high packing density is also relevant for studies on GUV-GUV adhesion ^69^ or while building multibody GUV tissues. ^70^

**Figure 3:**
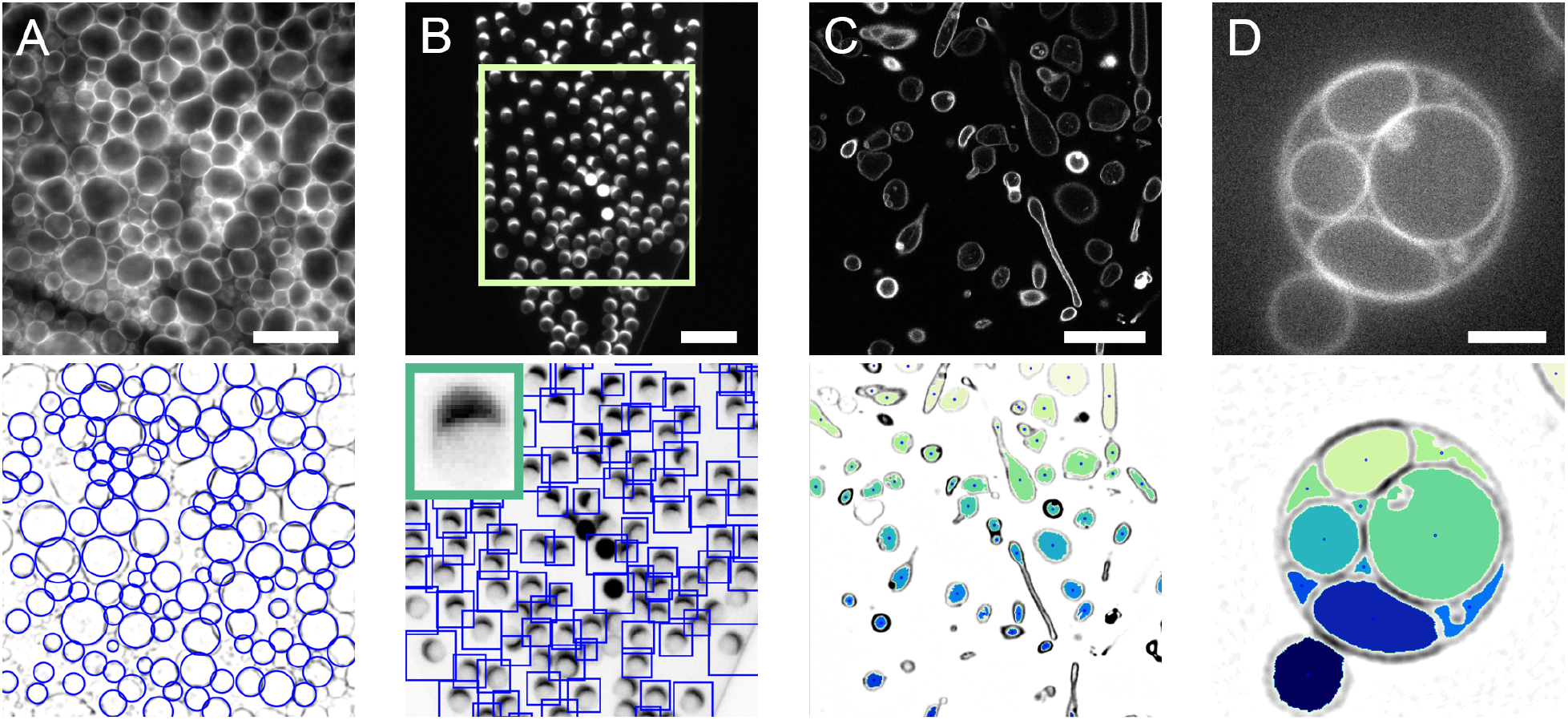
Applications of vesicle detection methods. Images at the top are the input fluorescence images, images at the bottom show the detection results. Contrast has been inverted for detection results to improve visualisation. (A) GUVs growing on top of a hydrogel following the gel-assisted swelling method are detected with CHT. Scale is 40*µm*. (B) Microfluidic production of GUVs imaged with a low magnification objective. MTM is employed to detect GUVs in the microfluidic channel. Produced vesicles contain a lipid-rich octanol pocket, visible as a bright cap. Scale is 100*µm*. Inset in bottom image is the template used for detection. (C) Encapsulation of stiff actin bundles in GUVs leads to deformation of the vesicles (data by F.C. Tsai, from ^40^). The non-spherical vesicles can be detected with FF detection. Scale is 20*µm*. (D) A GUV formed by gel-assisted swelling contains large internal vesicles. FF detection on this multivesicular GUV leads to detection of the compartments rather than detection of the enclosing GUV. Scale is 5*µm*.

As an alternative to the classical swelling methods, microfluidic vesicle production is becoming increasingly popular, with multiple new techniques being published yearly (reviewed in ref. ^71, 72^). Microfluidics offers superb control over vesicle formation, making it a powerful tool in the synthetic cell engineering field. Vesicles can be imaged *in situ* as they are being produced on-chip, using objectives with a large working distance with low magnification. This typically yields low-resolution images of vesicles. In line with MTM’s ability to detect out-of-focus vesicles (fig. 2H), MTM also proves to be suitable for GUV detection in low-resolution images of microfluidic GUV fabrication, as we demonstrate with an octanol-assisted liposome assembly (OLA) experiment^22^ (fig. 3B). Detection of vesicles on-chip enables users to extract GUV production rates and corresponding size distributions in microfluidic experiments. MTM detection does not require a sharp outline of the vesicle, but only a template that resembles the vesicles that need to be detected. Since the template can easily be picked from the image itself, MTM provides a versatile tool for vesicle detection even in low-resolution images. As alternative to MTM, CHT is also a useful detector in microfluidic experiments, since production by microfluidics often leads to spherical vesicles at high packing density with a narrow size distribution. ^57^

While CHT and MTM are both shape-sensitive detectors, FF can detect vesicles of any shape. Having a detector that does not rely on vesicle shape is valuable, as shape control and GUV deformation are essential aspects of synthetic cell engineering. ^9, 40, 73, 74^ We demonstrate the use of FF on GUVs deformed by encapsulated stiff actin bundles (fig. 3C). In this experiment, filamentous actin is co-encapsulated with the bundling protein fascin, resulting in the formation of actin bundles up to tens of micrometers long.^40^ Due to the high stiffness of these bundles, GUVs are deformed, resulting in elongated vesicles and actin-filled membrane protrusions. In fig. 3C, it can be seen that FF detects all vesicles irrespective of shape. In turn, detection by FF can be used as a starting point for vesicle deformation studies. Another application of the FF detector is found in the image segmentation of GUV internal compartments. Just like living cells contain numerous reactive compartments, GUVs can be compartmentalized to spatially separate cellular processes. ^75, 76^ Compartmentalization is becoming more popular in synthetic cell research, as distinct reaction environments are desired for reconstitution of increasingly complex processes. Due to its shape-insensitive detection, the FF method is suitable for detecting GUV compartments with random shapes and sizes as illustrated in fig. 3D. In this way, detection of compartments could be used to monitor internal activity of cellular processes.

Altogether, the three vesicle detection methods make DisGUVery useful for a wide range of synthetic cell research applications. Based on properties of the sample inspected, such as image resolution and vesicle shape, the desired detection method can be chosen (fig. 4).

**Figure 4:**
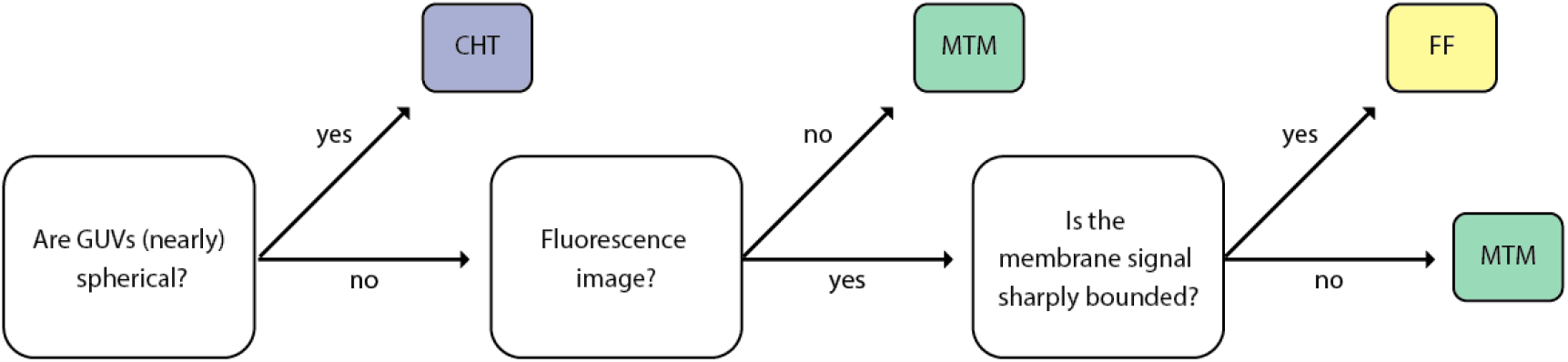
Decision tree for choosing one of DisGUVery’s detection modules based on sample and image properties.

### Membrane analysis

Membrane detection and analysis are important for a wide range of GUV studies, as they allow vesicle shape characterisation and quantitative analysis of the membrane fluorescence and/or membrane-binding proteins or other molecules. We have implemented two modules to perform membrane analysis on the detected vesicles: Refined Membrane Detection (RMD) and Basic Membrane Analysis (BMA). We have developed RMD to enable the tracking of the membrane contours, facilitating the capture and quantification of any global and local deformations. This method is based on a Canny edge detector that we have combined with a directional search algorithm to assign the detected edges to the inner and outer contours of the membrane (fig. 5A-C, contours shown in red). While the position of the detected contour can be affected by the choice of kernel size used with the edge detector (a predictable offset is introduced), the membrane position, taken as the midpoint between the inner and outer contour, will remain independent of the kernel size for confocal fluorescence images. Compared to RMD, BMA is a faster and simpler method to analyse the vesicle fluorescence signal, but at the expense of lower spatial accuracy. In BMA, a region of interest is created by a simple expansion of user-defined width around the boundaries of the detected vesicle. In case of CHT detection, this results in a circular ring as shown in fig. 5C.

**Figure 5:**
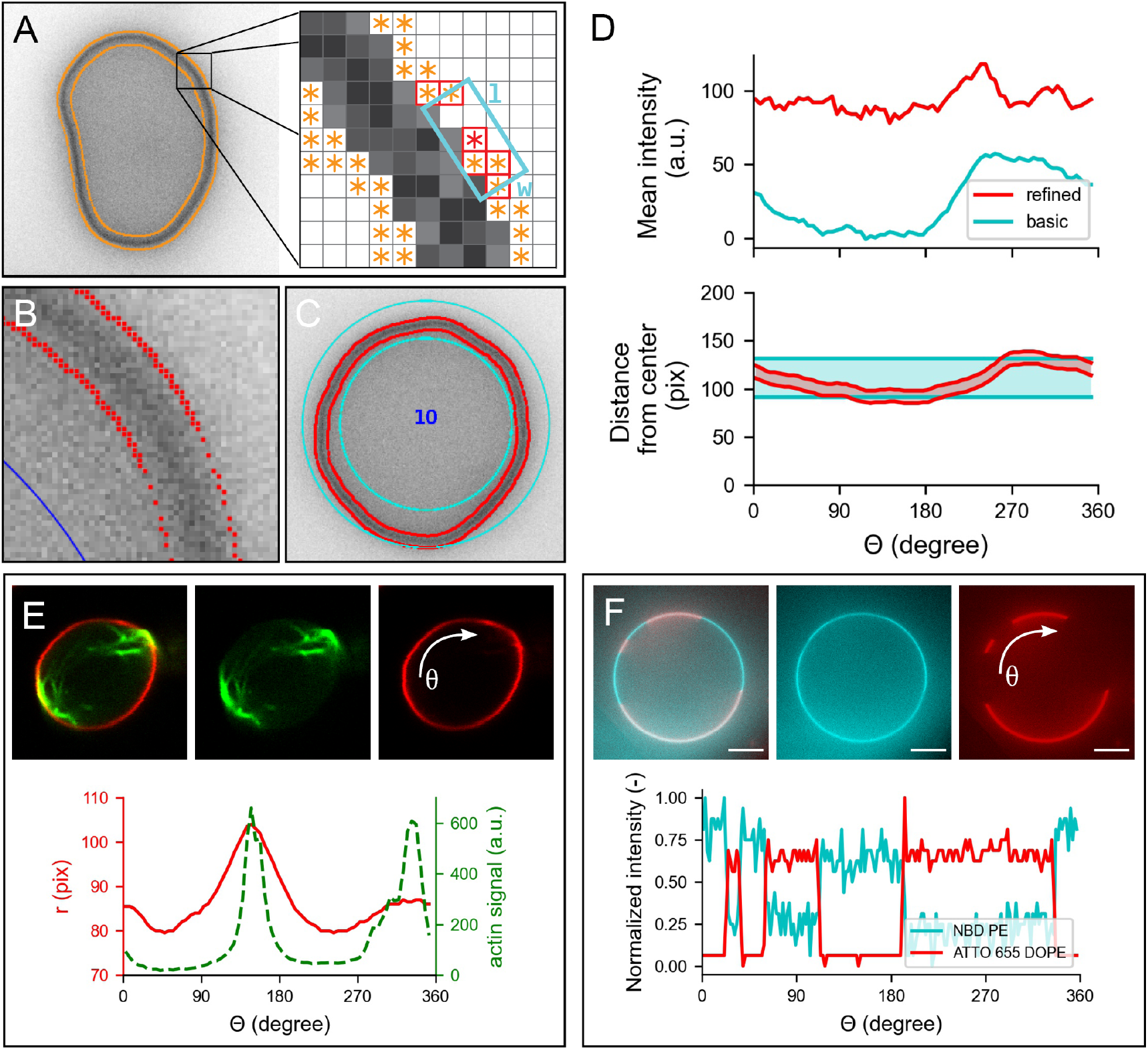
Membrane analysis by DisGUVery. (A) Schematic overview of the refined detection method. Left: epifluorescence image (inverted) of a GUV with all detected edges (orange). Right: edge points (orange) within the search box (cyan) of size *l* x *w* are connected (red border). (B) Zoom-in on membrane edges detected by refined detection (red), displayed on top of the inverted epifluorescence image of a GUV. (C) Segmentation of the membrane area as defined by basic membrane analysis (cyan) and refined membrane detection (red). (D) Angular profile of membrane properties from the vesicle in (C) extracted by basic membrane analysis (blue) and by refined membrane detection (red). Top: mean intensity per angular slice with an angular separation of 5°, and a ring width of 30 pixels for BMA. Bottom: radial distance to inner and outer boundary from the center of the vesicle. (E) Refined membrane detection on a non-spherical GUV deformed by actin bundles. Insets: composite confocal image of a GUV membrane (red) deformed by actin- fascin bundles (green) (data by F.C. Tsai, from Tsai et al. ^40^). Plot: angular profile of the membrane’s radial distance (red) and integrated actin intensity (green). (F) Basic membrane analysis of a phase-separated membrane containing DOPC:DPPC:cholesterol:NBD-DPPE:ATTO655-DOPE in a 31.8:48:20:0.1:0.1 molar ratio. Insets: image of vesicle labeled with NBD-DPPE (blue) and ATTO655DOPE (red). Plot: angular profile of both dyes extracted by basic membrane analysis, normalized to unity by subtraction of the minimum signal followed by division by the remaining maximum signal.

After membrane segmentation, either by contour detection in RMD or by defining a region of interest in BMA, it is possible to extract the angular and radial intensity profiles of the entire vesicle. The intensity profiles are calculated by creating angular or radial slices, of size Δ*θ* or Δ*r*, and computing the corresponding descriptive metrics for each slice, thus taking into account all intensity values of the detected vesicle and reducing the effects of discretization associated to single linear profile extraction. For the vesicle in fig. 5C, the angular intensity profile using the mean membrane intensity from each angular slice is shown in fig. 5D (top). Note that, although the trend of the mean intensity profiles is similar for RMD and BMA, the values differ greatly. This is a consequence of the wider segmentation ring of BMA (fig. 5D, bottom), which, when used to compute the mean intensity values, introduces the influence of the background signal, unlike the contained segmentation done by RMD. Furthermore, it can be seen that the BMA profile shows a fluorescence increase from *θ* = 250° to 300°, while this effect is much weaker for RMD. We attribute this apparent increase in fluorescence to the fact that the membrane shows an outward deformation around *θ* = 250° (fig. 5D, bottom), causing a larger part of the BMA slices starting from that angle to be filled with membrane as compared to other slices. Since in RMD the ROI always tightly confines the membrane, extraction of fluorescence intensity is much less sensitive to membrane shape. Dependent on the nature of the data and the required analysis, the choice of descriptive metrics can have a significant contribution of imaging artefacts or other sample related noise (see SI). For example, while both the integrated intensity and the mean intensity are influenced by the background signal, the latter will also depend on the number of pixels within the slice. Polydisperse samples, where the vesicle size has a large variation, will thus require a careful interpretation of the results and likely, a different metric to analyse the data as compared to more monodisperse samples.

To further illustrate the applicability of both methods in the quantitative characterisation of GUV membranes, we show how RMD can be used to analyse the membrane and content of a deformed vesicle (fig. 5E), while we use BMA for an example on phase-separated membranes (fig. 5F). In the first example, a GUV is deformed to a prolate shape by encapsulated filamentous actin that is bundled by the bundling agent fascin. ^40^ We used RMD to track the membrane contour position, and additionally, we extracted the angular profile for the average actin intensity from the RMD contour (fig. 5E bottom). The plot clearly shows two peaks in membrane position around angles 150 and 340°, which correspond with the peaks in actin intensity. If desired, the obtained contour coordinates can be exported to compute other shape descriptors of interest. In this way, membrane deformation by fluores-cent structures can be quantified in an automated way for vesicle populations, enabling an accessible and quantitative approach in GUV deformation studies. Besides its use in actin-mediated GUV deformation studies, ^40, 44, 77–79^ this analysis is also valuable in other studies on global vesicle shape deformation, for example by other proteins involved in cytokinesis such as the bacterial division proteins FtsZ^80^ and Min system, ^73^ by other membrane-binding proteins, ^9^ DNA origami, ^81^ by microfluidic traps, ^74, 82^ or by spontaneous membrane fluctuations. ^83^ Furthermore, RMD could be applied to characterize local membrane deformations, such as protrusions^84^ or nanotubes. ^28, 54^

In cases where vesicles are rather spherical and their shape well characterised, BMA is a useful tool to study the fluorescence signal of the GUV membrane. In fig. 5F, we show a GUV composed of a lipid mix of DOPC:DPPC:cholesterol:NBD-DPPE:ATTO655-DOPE in molar ratio 31.8:48:20:0.1:0.1. In this ratio, the lipids form two spatially separated phases: ^85^ a liquid-ordered phase containing mainly DPPC lipids and cholesterol, and a liquid-disordered phase containing mainly DOPC lipids. While NBD-DPPE partitions preferentially into the liquid-ordered phase (blue), ATTO655-DOPE accumulates in the liquid-disordered phase (red). Using CHT detection followed by BMA, we obtained the angular profiles of both membrane dyes. The intensity profiles indeed clearly reveal that both dyes have a preferential presence in either one of the two phases. While here we show the example analysis for one single GUV, we would like to stress that BMA performs membrane analysis at high computation speed, enabling the analysis of membrane fluorescence for hundreds of vesicles within minutes. Next to lipid-lipid phase separation studies, BMA could be used for membrane quenching experiments,^86^ to probe the homogeneity of a reconstituted actin cortex, ^30^ or for analysis of spectral images in lipid packing studies using polarity-sensitive probes. ^87^

### Population analysis

So far, we have demonstrated DisGUVery’s working principles and the performance of detection and membrane analysis on single vesicles or single images. However, for the analysis of GUV experiments it is often desired to analyse large numbers or time-lapse series of vesicles. We implemented a batch-processing option that allows for the semi-automated analysis of multiple images, making population characterisation on large data sets accessible and enabling easy identification of statistical differences. We illustrate the potential of the batch-processing feature with two quantitative analyses: binding of small vesicles to GUVs using membrane-anchored oligonucleotides, and the encapsulation of a fluorescent protein inside GUVs.

In the first case, we utilise the membrane analysis module on a population of vesicles where we bound large unilamellar vesicles (LUVs) to GUVs by using membrane-anchored oligonucleotides (fig. 6A-E).^88, 89^ While GUVs have diameters of tens of microns (fig. 6A), the LUVs used in this study have a diameter of approximately 200 nm, close to the size of the diffraction limit. To generate specific binding between GUVs and LUVs, we incorporated one type of single-stranded DNA (ssDNA) in the GUVs, and the complementary ssDNA in LUVs. Here, we set out to test if the extent of LUV-GUV binding could be regulated by varying the DNA concentration. Therefore, we incubated vesicles either with 0.5*µM* DNA, 1*µM* DNA or no DNA at all prior to mixing LUVs with GUVs. LUVs were doped with a fluorescently tagged phospholipid for visualization and quantification. To allow vesicle detection that is not biased by LUV-binding, we independently labeled GUV membranes with another fluorescent phospholipid. When both types of vesicles were incubated with 1*µM* of the complementary DNA strands prior to mixing, LUVs clearly localized on the GUV membranes (fig. 6B, fig. S3B), while we observed no colocalization in absence of DNA(fig. S3A). To quantify LUV binding, we first detected GUVs in the Atto488 channel using CHT detection (fig. 6C). Membrane fluorescence was analyzed using the basic membrane analysis because it is computationally light and our analysis did not demand a high spatial accuracy (fig. 6D). We chose a large (50 pixel) ring width to be able to extract membrane fluorescence also from non-spherical vesicles that were naturally present in the sample. While the software exports different intensity metrics from the angular slices, we performed our analysis using the intensity maximum per slice to minimize the effect of the background signal (see SI). Given that the maximum is sensitive to fluorescence outliers, for example caused by bright membrane structures or touching vesicles, we finally take the median of all angular maxima to represent the vesicle average. To correct for background intensity, we subtract the radial intensity average just outside the vesicle from the vesicle-average LUV intensity. In this way, we analyzed over 1000 GUVs in 50 different images. The results are plotted in fig. 6E. In absence of DNA, LUVs do not bind to GUVs, in line with what is seen the image (fig. S3). Upon DNA addition, membrane analysis shows a clear increase in membrane localization of LUVs. Furthermore, quantitative membrane analysis reveals that the LUV intensity is significantly higher when using 1 *µM* DNA than 0.5 *µM* DNA (p*<<*0.001). The data in fig. 6E underlines why population statistics can be essential for analyzing GUV data sets. While vesicles with similar LUV intensity exist in both populations, a statistical difference between the two populations can only be proven when a large number of vesicles is analyzed. In this way, high-throughput membrane analysis helps to quantitatively investigate the effect of experimental parameters on GUV membrane studies.

**Figure 6:**
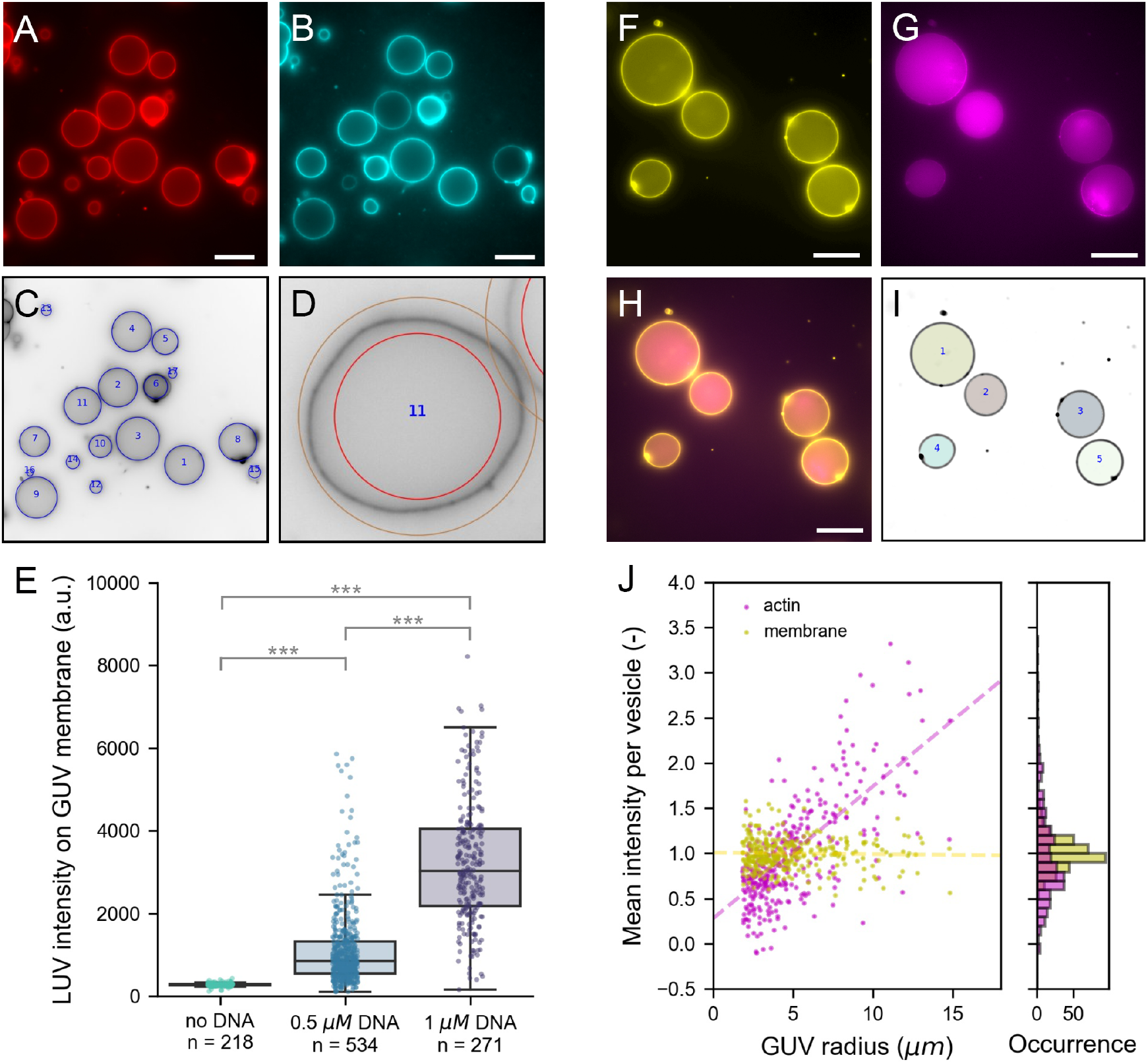
Population analysis. (A-E) Analysis of LUVs binding via membrane-anchored oligonucleotides to GUV membranes. (A) Atto 488 DOPE-labeled GUVs produced by gel-assisted swelling. (B) Atto 655 DOPE-labeled LUVs localize on GUV membranes when both are incubated with 1 *µM* cholesterol-DNA. (C) CHT detection in the Atto488-channel (inverted contrast). Detected vesicles are indicated with blue circles. (C) Example of the detection ring of 50 pixels width used for basic membrane analysis. (E) Bar plot of LUV intensity on the GUV membrane at different DNA concentrations. Each point represents the LUV intensity on an individual vesicle. *** indicates statistically significant difference with p *<* 0.001. (F-J) Analysis of fluorescent monomeric actin encapsulated in GUVs using cDICE. (F) DOPC GUVs labeled with 0.1% (mol/mol) 18:1 Cy5 PE. (G) Encapsulated actin of which 10% is labeled with Alexa 488. (H) Composite image of membrane and actin. (I) Results of FF detection. Masks represent detected vesicles. (J) Mean intensity normalised by population average of actin (magenta) and membrane (yellow) plotted against the GUV radius (left) and shown in a histogram (right). Dashed lines in the scatter plot are linear regression results for actin (magenta, slope is 0.15) and membrane (yellow, slope is 0.00). All images are epifluorescence images. Scale bar is 20 *µm* in all images.

In the second example of population analysis, we perform an encapsulation analysis using the Encapsulation Analysis module in DisGUVery. Encapsulation of molecules, proteins, vesicles and even living cells inside GUVs is becoming more and more important as GUV-based reconstitution experiments are increasing in complexity.^90, 91^ Besides controlling which type of molecules end up in the GUVs, also their concentration and stoichiometry often need to be regulated in order for them to function properly. It is essential to evaluate the quality of encapsulation, as this varies substantially between experiments, depending strongly on the way the GUVs are produced as well as on the molecule that needs to be encapsulated. ^12, 25, 92^ When the encapsulated molecule can be visualized with fluorescence microscopy, the encapsulation efficiency can be determined as the distribution of internal fluorescence of the encapsulated molecule across the GUV population. To demonstrate this, we encapsulated monomeric actin in GUVs using the continuous Droplet Interface Crossing Encapsulation (cDICE) technique ^24^ following the protocol outlined by Van de Cauter et al. ^25^ In this experiment, 10% of the actin monomers were labeled with Alexa 488 to allow for fluorescence visualization. Vesicles were imaged in epifluorescence microscopy to capture the signal of the entire vesicle volume in a single frame. From fig. 6F-H, it can be seen that the actin signal is strongly enhanced inside the GUVs as compared to the outer solution, and that the observed fluorescence varies among vesicles. To quantify the encapsulation efficiency, we first detected vesicles based on the membrane signal by means of the FF detection method (fig. 6I). The advantage of using FF detection is that detected masks directly match the projected shape of the vesicle lumen, independent of vesicle shape and size. Using the output of the FF detection, we extracted the mean intensity of both the actin and the membrane signal for each vesicle. Furthermore, in each image, we determined the background signal for each imaging channel by taking the mode of the intensity histogram. Background signals were subtracted from the mean intensity per vesicle to finally yield the corrected mean intensity per vesicle. In total, we analyzed 329 vesicles in 22 images of one preparation. In fig. 6J we show the distribution of the corrected mean intensities for actin and the membrane. Note that in the epifluorescence imaging mode, the fluorescence emission from the entire focal volume is projected onto the imaging plane. Since the focal depth of the system is larger than the vesicle size, we expect a clear dependency on the vesicle size for any fluorescent molecule distributed in the volume of the GUV. In contrast, the membrane fluorescence signal is localized in the surface area of the vesicle, meaning that always the same volume of fluorescent membrane probes is projected onto the focal plane, independent of vesicle size. As a result, when we plot the mean intensity as a function of radius, we can easily distinguish encapsulated proteins from those that localize on the membrane. This offers an alternative route to probe fluorescence localization despite the lower depth-sectioning of epifluorescence imaging compared to confocal microscopy, enabling faster image acquisition and facilitating the screening of large datasets. We observe the expected linear trend for the intensity of encapsulated actin as a function of GUV radius (fig. 6J, dashed line) indicating that actin is distributed through the vesicle volume. The membrane mean fluorescence, on the other hand, shows no dependency on the vesicle radius, confirming that the fluorescent probe is membrane-bound. Furthermore, the mean intensity spread within the same vesicle size is larger for actin than it is for the membrane, reflecting the variability from protein encapsulation across vesicles. This analysis yields a relative measure of variations in encapsulation efficiency among GUVs, which, once combined with a calibration, could be used to evaluate absolute concentrations inside GUVs.

## Conclusion

Giant Unilamellar Vesicles have become a widely used system for research in biophysics and synthetic biology. As the versatility and complexity of applications grow, and in concert the number of GUV formation methods, it becomes increasingly important to perform rigorous and standardised quantitative analyses. Here, we presented DisGUVery, an open-source software that we have developed for the high-throughput detection and analysis of GUVs in a wide range of microscopy images.

Since the detection of GUVs is the first step in any type of analysis, we have done an in-depth characterisation of the object detection algorithms that we have adapted and implemented. Our results show that each detector can be used as a filter for specific vesicle types, and that we are able to overcome the influence of imaging source by careful selection of the detector. By testing and demonstrating detection in a broad range of typical GUV samples, we show that DisGUVery fits in with many areas of GUV research. So far, the simplicity of GUVs combined with our hands-on experience in GUV research have allowed us to develop lightweight algorithms with good detection performance. However, we note that with increased morphological complexity, it might be necessary to use more complex detectors, such as supervised machine learning. Even then, our software can serve as an accessible basis for generating training data sets for machine learning, thanks to the automated high-throughput segmentation algorithms.

As many GUV studies rely on shape and fluorescence of the membrane, we implemented a set of tools for membrane segmentation, which can be chosen depending on the spatial accuracy needed. Notably, we developed a membrane contour tracking method by coupling an edge detector with a directional search algorithm that takes advantage of the unique intensity profile of the membrane fluorescence. Furthermore, we showed how a contained membrane segmentation can easily identify local deformations and be less influenced by the background signal, when compared to a user-defined ROI that segments the membrane. Nevertheless, we illustrated how even a basic segmentation, in combination with high-throughput analysis, can identify statistical differences between GUV populations. Although we focused here on the intensity of the membrane, note that DisGUVery also allows to obtain the angular and radial intensity profiles of any imaging channel, allowing the user to study spatial distribution of encapsulated content. Altogether, the membrane analysis modules can be used to extract a wide range of vesicle properties, including GUV shape, internal fluorescence, membrane localisation of fluorescent proteins, or formation of internal structures. As such, the software can be used for all sorts of assays, such as membrane permeabilization studies, reconstitution of cytoskeletal networks, microfluidic vesicle production, GUV deformation studies, or membrane fusion assays. Although DisGUVery has been developed originally for detection and analysis of vesicles, the software might be equally useful for data analysis in other research domains involving similar types of microscopy data, such as colloidal and interfacial science.

Concluding, DisGUVery offers an accessible way to perform fast but thorough quantitative analysis of GUV microscopy images. By combining versatile vesicle detection and analysis algorithms, the software can robustly be employed for any type of GUV research. This makes DisGUVery a powerful tool that will help the field to progress towards a more quantitative, population-based research.

## Materials and methods

### Software algorithms

DisGUVery will be available shortly on an open-source repository.

### Image preprocessing

DisGUVery includes two image preprocessing options to aid vesicle detection: smoothing and membrane enhancement. Image smoothing is performed by convolution of the original image with a 2D Gaussian function of user-defined kernel size. Membrane enhancement is based on the subtraction of a second smoothed image (the subtraction image) from the first smoothed image. To this end, the user specifies a kernel size (typically 10-20 times larger than the kernel used for smoothing the image) to create the subtraction image also based on Gaussian smoothing, which is used to eliminate large scale intensity variations. The subtraction image is then subtracted from the smoothed image and all pixels with negative intensity values are set to zero, creating the membrane enhanced image.

### Circular Hough Transform detection

We implemented a Circular Hough Transform detection method ^62, 93^ based on the HoughCircles function from the OpenCV package. ^94^ The procedure consists of two steps: edge detection and circle detection. First, edges are detected in an image of size *x × y* using a classical Canny edge detector. The user can pass an edge threshold to the detector to filter out low quality edges. Using the detected edges, the Circular Hough Transform is applied for circle detection. The method is well described in ref. ^93^ In short, circle detection is done via a ‘voting’ procedure by voting in the Hough parameter space. To this end, the user passes the minimum and maximum radius for detection, after which the range is discretized into *N* radii. A three-dimensional accumulator array of the size (*x, y, N*) is created to record votes, where high numbers of votes represent the circle centers. Initially, the value of all cells in the accumulator matrix is set to zero, after which the voting procedure is done as follows. For each edge point (*i, j*) in the original space, a circle is formulated in the Hough parameter space centered at (*i, j*) with a certain radius *r*. For each point (*a, b*) in parameter space that the circle passes through, i.e. that fulfills (*i − a*)^2^ + (*j − b*)^2^ = *r*^2^, the voting number is increased by one in the accumulator matrix in the corresponding point (*a, b, r*). In this way, the accumulator value increases for points where circles in Hough parameter space intersect, which correspond to the circle centers in original space. The procedure is repeated for all radii *r*. Finally, the local maxima in the matrix are detected to yield the circle center coordinates (*x_c_, y_c_*) and their radius *r_v_* . By passing a Hough threshold to the detector, the user can selectively detect circles with the highest number of votes, corresponding to the highest quality circles. In addition, the user inputs a minimum radius between circles, which is used to prevent detection of multiple local maxima per GUV. Detected circles are drawn in the main display. It should be noted that detected radii and vesicle centers are approximations because of two reasons. One, the possible radii are discrete values, following from a discretization of the radii between *r_min_* and *r_max_* into *N* values. Two, the membrane signal in the input image represents an intensity maximum, meaning it provides two edges (an inner and an outer) for the Canny edge detector. Since both edges are close together, voting results from both edges might overlap, introducing an inaccuracy in the accumulator values. Refinement of spatial detection of GUV center and image can be done with Refined Membrane Detection.

### Multiscale Template Matching detection

DisGUVery’s Multiscale Template Matching detection is based on the OpenCV ^94^ function matchTemplate with an added template re-scaling option to enable size-invariant detection. First, a template needs to be selected: a template can be either defined manually from the input image, or a template image can be loaded. For template detection with a template of size *w× h* in an image of size *x× y*, the function compares overlapped patches of size *w× h* of the image against the template. By default, the comparison method cv2.TM_CCOEFF_NORMED is applied where a high matching coefficient results in a high comparison value. A map of the comparison results is created by storing the comparison results in an array of size (*x−w*+1)*×*(*y−h*+1). Object detection is then performed by finding the local maxima in the comparison map. The user can pass a matching threshold to retain only high-quality results.

Multiscale template matching is performed by resizing the template with a scaling factor *a* using OpenCV function resize to templates of size *aw × ah*. Scaling factors are computed by the software from the user inputs minimum rescaling factor, maximum rescaling factor, and number of scales. The output generated by the software are the object locations *x_c_, y_c_* and the bounding boxes of size *aw × ah*.

### Floodfill detection

Vesicle detection by Floodfill is implemented as follows. It is highly recommended to start with membrane enhancement for FF detection, as detection relies on absolute intensity differences. The most important step for Floodfill detection is the binarization of the image. The image is first thresholded with a user-defined relative threshold value. The absolute threshold is computed by multiplying the input value with the median of all positive pixels (intensity higher than 0). Then, the absolute threshold is used to binarize the image, the result of which can be inspected in the GUI. Here, foreground pixels or membrane pixels have intensity value 1 (white), while background pixels have value 0 (black). DisGUVery’s FF detection works by flooding background pixels in the outer solution which are connected using the FloodFill function of the OpenCV package. When pixels are flooded, their intensity is set to a different number. In this way, all background pixels in the exterior solution get a different value from background pixels in the GUV lumen and thus can the GUV interiors be distinguished. Flooding is started from a single point, the seed point, which must be located in the outer solution. After flooding, a binary mask of the size of the image is created where pixels belonging to the lumen of all GUVs are set to 1. The lumen masks will naturally contain holes, due to interior membrane structures or noise. These holes are filled by performing a second floodfilling step on the binary mask, flooding all pixels in the image except for the GUV interiors and the holes in the GUV masks. The filled GUV masks can then be collected by selecting all pixels with intensity value 0 and 1 and are stored in a second binary mask. All connected pixels are detected as a single GUV, after which objects smaller than the user input minimal area are discarded, and each separate GUV is labeled as a different object. Finally, DisGUVery creates as output the *x*, *y* matrix containing the labeled objects.

### Basic Membrane Analysis

Basic Membrane Analysis can be performed after vesicle detection. First, for each detected vesicle a region of interest (ROI) is created around the membrane based on the detection results. The ROI can be inspected in the image, after which angular and radially integrated intensity profiles are extracted from the ROI. The region of interest with a selected width *w* is defined differently dependent on the GUV detection method used. For a vesicle detected by CHT with radius *r*, two concentric circles are drawn from the vesicle center with *r_in_* = *r−w*/2 and *r_out_* = *r*+*w*/2 that together form a ring-like ROI. To obtain the angular intensity profile, the area confined by the inner and outer edge is divided into *n_a_* angular slices calculated from the user input angular width per slice Δ*θ* (°). From each slice, various region properties for fluorescence intensity (minimum, maximum, average, background-corrected average and sum) are obtained. Furthermore, the radial distance of the ROI inner and outer edge, *r_in_*(*θ*) and *r_out_*(*θ*), respectively, can be extracted. Similar to extracting the angular intensity profile, the radial intensity profile is obtained by specifying the radial integration width Δ*r* (pixel), used to calculate the number of radial rings *n_r_*. Integrating over all angles, the software then extracts region properties for all radial rings from the vesicle centre *r*_0_ = 0 to *r_out_*(*θ*). For each radial ring, various intensity metrics are obtained (minimum, maximum, average, background-corrected average and sum).

### Refined Membrane Detection

The Refined Membrane Detection method consists of two steps: first all edges in the image are detected (the edge tracking step), then edge points are chained together and assigned to either the outer or the inner edge of a GUV membrane (the chaining step). Edges are tracked with a modified Canny edge detection algorithm with the wavelet transform (WT) as outlined in ref. ^95^ By using the first derivative of a Gaussian in the WT, the 2D smoothed gradient of the image is computed. ^96^ In addition, for each point in the image the amplitude of the gradient (the WT modulus) and its direction (the WT argument) are obtained. These are passed to a modified Canny edge detector^97^ in order to compute the edges. At each image coordinate, a point is considered to be an edge if the wavelet transform modulus is a local maximum, or the intensity in that point has the steepest gradient, when compared to its neighbouring pixels. The comparison is made with the pixels that follow the argument of the wavelet transform at that point, that is, in the direction of the steepest gradient which is across the membrane. To distinguish ‘true’ edges from falsely detected edges, points are chained together and a double hysteresis algorithm is applied to connect weak with strong edges. ^97^ This process finally results in an edge mask that contains all the true edges of the entire image. For the second step, we developed a directional search algorithm to chain edge points and to assign them to either the inner or the outer edge of the membrane. The search occurs at each detected edge point, chosen at random, by chaining the neighbouring points contained within a defined search box centred around the point of interest and typically of aspect ratio *>* 2:1 (length: width). The orientation of the box is determined as orthogonal to the argument of the wavelet transform at the chosen point. Given that the WT argument will follow the direction where the gradient is steeper, thus across the membrane, its orthogonal direction will likely follow the points along the membrane. Lastly, all chains are measured by the number of points within, and the two longest chains will be assigned to the outer and inner edge of the membrane (fig. 5B). This directional search with a bounding box allows to chain points together without the need of them being connected. Furthermore, it enables the user to distinguish the enclosing GUV membrane, which is the membrane separating the inner from the outer solution, from internal membranes and secondary membrane structures such as tubes. Dependent on the membrane appearance, for example its thickness in the image, or the presence of secondary membrane structures, the user can define the length and width of the search box to fine-tune the tracking results. Finally, two lists of *x* and *y* coordinates are produced for the inner and outer edge of the GUV membrane. By calculating the centre of mass of the detected contours, the vesicle centre is refined. The angular profiles of radial distance from the vesicle centre to the inner and outer edge are calculated and can be exported. Furthermore, an ROI is defined between the two membrane edges, which can be used to extract angular and radial fluorescence profiles such as described above for BMA.

### Encapsulation analysis

The Encapsulation Analysis module uses masks of GUV interiors to collect all fluorescence intensities from under those masks. The user has various options to create the masks. First, the user can directly use the results from one of the three GUV detection methods as a mask. This yields circular masks for CHT, rectangular masks for MTM, and free shape masks for FF detection. Depending on the detection method chosen and the spatial accuracy required, the user can choose to perform an additional refinement step to refine the shape of the mask. In this refinement step, the image is first segmented using rectangular bounding boxes defined by the vesicle detection results. Then, for each GUV in its bounding box, FF detection is applied as described above to yield masks with fitting shape.

## Experimental data

### Chemicals and proteins

From Avanti Polar Lipids we obtained the lipids L-*α*-phosphatidylcholine (eggPC), 1,2- dioleoyl-sn-glycero-3-phosphocholine (DOPC), 1,2-distearoyl-sn-glycero-3-phospho-ethanolamine-N-[methoxy(polyethylene glycol)-2000] (PEG2000-DOPE), 1,2-dipalmitoyl-sn-glycero-3-phosphocholine (DPPC), 1,2-dioleoyl-sn-glycero-3-phosphoethanolamine-N-(Cyanine 5) (Cy5-DOPE), 1,2-dipalmitoyl-sn-glycero-3-phosphoethanolamine-N-(7-nitro-2-1,3-benzoxa diazol-4-yl) (NBD-DPPE) and 1,2-dioleoyl-sn-glycero-3-phosphoethanolamine-N-(lissamine rhodamine B sulfonyl) (Rhodamine-DOPE). The lipids ATTO 488 DOPE and ATTO 655 DOPE were obtained from ATTO-TEC GmbH (Siegen, Germany). All lipids were stored in chloroform at -20 °C under argon. The chemicals D-(+)-glucose, sucrose, Tris-HCl, KCl, 1-octanol, glycerol, Poloxamer 188, cholesterol, dithiothreitol (DTT), protocatechuic acid (PCA) and the proteins protocatechuate dioxygenase (PCD) and *β*-casein were obtained from Sigma Aldrich. For gel-swelling, we used poly-vinyl alcohol (PVA) of 145 kDa, 98% hydrolysed, obtained from VWR, Amsterdam, the Netherlands. Actin for the encapsulation experiments was purified in house as described in ref. ^98^ Alexa-488 labeling of actin was done in house following ref. ^99^

### GUV imaging

The protocols for GUV formation are described below. Depending on the formation method, produced GUVs were transferred to one of two types of imaging chambers: small 20*µL* wells assembled by using silicone gaskets, or 150*µL* wells made by using PCR-tubes. The small wells were used for samples of high GUV density, such as gel-swelling experiments, while the large wells were used to image GUVs produced at lower concentrations such as eDICE. For the preparation of small imaging wells, we first rinsed a 24x50*mm* cover glass (No. 1.5H, Thorlabs) with ethanol, water and ethanol, and then blow-dried it with nitrogen. Then, an 8-well silicone spacer (6 mm diameter x 1 mm depth, CultureWell^TM^, Grace Bio-Labs) was pre-wetted with isopropanol, dried with nitrogen gas and placed on top of the glass. To each well, we added 15 *µL* of *β*-casein solution (1 mg/mL *β*-casein, 10 mM Tris-HCl pH 7.4) and let it rest to passivate the glass surface against membrane adhesion. After 15 minutes, the solution was removed with a tissue and the chambers were blow-dried with nitrogen gas. We added to each well first 15 *µL* of an isotonic glucose solution and then 5 *µL* of vesicle solution. Due to their higher density, vesicles sunk to the bottom of the chamber, which facilitated imaging. The chamber was then closed from the top with a glass slide (1mm thickness, Thermo Scientific) to prevent solvent evaporation and minimize flow in the sample. Large imaging chambers were made by gluing a PCR tube with a cut bottom upside down on a 24x50 mm coverslip that had been cleaned as described above. The chamber was passivated by treating it for 15 minutes with a 1 mg/mL *β*-casein solution in milliQ water containing 10*mM* Tris-HCl at pH 7.4, after which the chamber was rinsed with milliQ water and blow-dried. Then, 150*µL* of vesicle sample was added and the chamber was closed by placing the lid of the PCR tube on top.

All GUV images (except fig. 3C and fig. 5E, as described below) analyzed in this work were obtained using an inverted microscope (Nikon Ti Eclipse) with a digital CMOS camera (Orca Flash 4.0). The different objectives and imaging settings used for the respective experiments are specified below.

### GUV formation protocols

The vesicles that were used to illustrate DisGUVery’s workflow (fig. 1A), to test detection (fig. 2), the high density GUVs growing on top of a hydrogel (fig. 3A) and the compartmentalized vesicle (fig. 3D) have been produced by PVA(poly-vinyl alcohol)-assisted swelling following ref. ^19^ with minor modifications. In short, glass coverslips (24 x 24 mm, Menzel Glaser) were rinsed with water and ethanol, blow-dried with nitrogen, and plasma-treated for 30 seconds to create a clean and reactive surface. Then, 100*µL* of a viscous 5% (w/v) PVA solution was spread over the coverslip to create a thin layer. The coverslip was baked in the oven at 50 °C for 30 minutes to form a gel. After baking, 10*µL* lipids in chloroform at a total lipid concentration of 1*mg/mL* was spread over the gel using a Hamilton syringe. Typically, membranes consisted of 99.9% EggPC lipids and 0.01% fluorescent ATTO 655 DOPE. GUVs were swollen for 60 minutes in a swelling solution containing 200 mOsm sucrose and 10 mM Tris-HCl at pH 7.4. After formation, GUVs were harvested and transferred to a small imaging well (described above) containing an isotonic glucose solution (200 mOsm glucose, 10 mM Tris-HCl at pH 7.4). The images in fig. 1, fig. 2 and fig. 3D were taken using a 100x oil immersion objective with a phase ring (NA 1.45, Ph 3, Nikon) at an exposure time of 100*ms* for all imaging methods used. For epifluorescence imaging, the sample was illuminated with monochromatic LED light of 640*nm* (Lumencor Spectra Pad X), at 25% of the maximum power. Confocal images were gathered on the same imaging set-up using a spinning disk confocal (Crest X-light) with pinhole size 70*µm* and illumination at 75% of full intensity. Phase contrast images were acquired with the microscope’s DIA illuminator switched on at a voltage of 10.5*V* and using the corresponding phase mask in the microscope’s condenser. For the image shown in fig. 3A, the coverslip with PVA gel and dried lipids on the microscope. The image was taken in epifluorescence mode using a 60x long working distance water immersion objective (CFI Plan Apochromat VS 60x WI, Nikon) 10 minutes after addition of the swelling buffer.

Actin-deformed GUVs shown in fig. 3C and also in fig. 5E were produced by F.C. Tsai as described in detail in ref. ^40^ In short, GUVs of a lipid composition of DOPC:Rhodamine-DOPE:PEG2000DPPE in a molar ratio of 94.8:0.2:5 were produced by gel-assisted swelling on top of an agarose gel. Actin was encapsulated by adding it to the swelling solution at a concentration of 12*µM*, and at a 5:1 molar ratio with respect to fascin. 20-30 mol% of the actin was labeled with Alexa488 to allow fluorescence visualization. After formation, GUVs were harvested and imaged by confocal fluorescence microscopy. Images were taken with a Nikon Eclipse Ti inverted microscope equipped with a Nikon C1 confocal scanhead, a 100x NA1.4 Plan Apo oil immersion objective and lasers with wavelengths 488 nm and 543 nm.

Microfluidic vesicle production (fig. 3B) was done with the octanol-assisted liposome assembly (OLA) technique following ref. ^22^ Lipids were used in a composition of DOPC: Rhodamine DOPE in molar ratio 99.5:0.5. The inner aqueous solution consisted of 5% (v/v) glycerol in milliQ water, and the outer solution of 15 % (v/v) glycerol and 5% (w/v) Poloxamer 188. GUVs were imaged directly on-chip in the post-formation channel with a 10x air objective (Plan Fluor, NA 0.3, Nikon).

Phase-separated GUVs (fig. 5) were produced by gel-assisted swelling as described above, but with minor modifications. Lipids were dried in a mixture of DOPC:DPPC:cholesterol: NBD-DPPE:ATTO655-DOPE in molar ratio 31.8:48:20:0.1:0.1. In addition, swelling was done in a 37°C room to be above the membrane transition temperature, and thus to ensure proper mixing of lipids during formation. GUVs were imaged in the small imaging chambers. DNA-mediated vesicle binding was performed following ref. ^88^ and ref. ^89^ . GUVs with a membrane composed of DOPC:ATTO 488 DOPE in molar ratio 99.5:0.5 were produced by gel-assisted swelling as described above in a solution containing 100*mOsm* sucrose, 100*mM* KCl and 10*mM* Tris-HCl at pH 7.4. LUVs were produced by adding lipids in chloroform to a Pyrex glass tube, in a lipid composition of DOPC:ATTO 655 DOPE as 99.95:0.05 (mol/mol). After drying lipids for 1 hour in a vacuum desiccator, the dried film was resuspended by vortexing for 2 minutes in a solution containing 100*mMKCl* and 10*mM* Tris-HCl at pH 7.4 to a final lipid concentration of 0.5*mg/mL*. To produce 200*nm* LUVs, the suspension was extruded (Mini Extruder, Avanti Polar Lipids Inc.) 21 times over a polycarbonate membrane with pore seize 200*nm* (Nuclepore, Whatman). To introduce specific binding between LUVs and GUVs, we used two complementary DNA strands (DNA1 and DNA1*^!^*) with a cholesterol moiety for membrane anchoring^88^ (biomers.net, Ulm, Germany). The DNA strands were tagged with cholesterol on opposite ends to allow antiparallel binding, as is typically used for DNA-mediated membrane fusion assays.^100^ The full sequences were taken from ref. ^89^ and read:

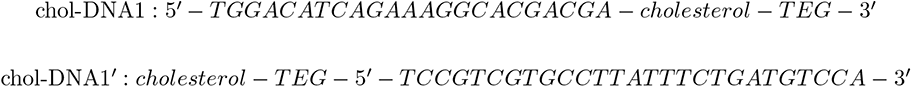

Note that the sequences do not fully overlap, which results from an error in the original publication (Y. Dreher, personal communication, 2021). For one hour, LUVs were incubated with 1*µM* of chol-DNA1, and GUVs with 1*µM* of chol-DNA1*^!^*. After DNA incbuation, GUVs and LUVs were mixed, left to bind for one hour, and finally imaged in the PCR tube imaging chamber in a solution containing 100*mOsm* glucose, 100*mMKCl* and 10*mM* Tris-HCl at pH 7.4. Images were taken in epifluorescence mode using a 100x oil immersion objective (CFI Plan Apochromat VC 100x oil, NA 1.40, Nikon) with an exposure time of 100 ms at 508 nm and 640 nm to image LUVs and GUVs, respectively.

Vesicles containing monomeric actin were produced as in ref. ^25^ . Lipids were mixed in a molar ratio of DOPC:PEG2000-DOPE:Cy5-DOPE 99.89:0.01:0.1. We encapsulated 4.4 *µM* actin in G-buffer of which 10% was labeled with Alexa-488 to allow fluorescence visualisation. The encapsulated solution also included 1*mM* dithiothreitol (DTT), 1*µM* protocatechuic acid (PCA) and 1*µM* protocatechuate dioxygenase (PCD). Epifluorescence images were taken with a 100x oil immersion objective (CFI Plan Apochromat VC 100x oil, NA 1.40, Nikon) at a wavelength 640 nm with 10% and 100 ms exposure time, and at 470 nm with 20% and 500 ms exposure time to visualise the vesicle membrane and actin content, respectively.

## Acknowledgements

We thank Martin Depken for useful discussions on evaluation of vesicle detection performance, Britta Bor, Gerard Castro Linares and Lucia Baldauf for performance evaluation of vesicle detection, Tom Aarts and Lucia Baldauf for images on DNA-mediated vesicle binding and Jeffrey den Haan for protein purification and labeling. This project was partly funded by ‘BaSyC - Building a Synthetic Cell’ Gravitation grant of the Netherlands Ministry of Education, Culture and Science (OCW) and the Netherlands Organization for Scientific Research (NWO).

