## Supporting Information for "DisGUVery: a versatile open-source software for high-throughput image analysis of Giant Unilamellar Vesicles"

### Membrane analysis metrics

DisGUVery's membrane analysis modules Refined Membrane Detection (RMD) and Basic Membrane Analysis (BMA) support the use of different metrics to extract membrane fluorescence. This section should serve as a guide to choose the right metric according to the imaging source and research goal.

#### One-dimensional intensity profile

We consider a lipid bilayer membrane dyed with a fluorescent lipid. Since the membrane is thinner than the diffraction limit, the fluorescence intensity profile is given by the point spread function (PSF). We assume the PSF to be a Gaussian function, for a one-dimensional signal given by:

$$I(x) = Ae^{-\frac{(x-r)^2}{2\sigma^2}} \tag{2}$$

where $x$ is the radial distance with respect to the GUV centre, $r$ is the GUV radius, $A$ is the amplitude of the signal and $\sigma$ is the standard deviation, or the width of the point spread function. $\sigma$ can be determined experimentally by fitting a Gaussian curve to the one-dimensional data.

In practice, fluorescence imaging is subjected to various sorts of noise (see fig. S4A). First, there is the camera read-out noise which contributes to a random noise in each pixel, $I_{cam}(x)$. Second, there is noise caused by out-of-focus fluorescence, ambient light and excitation light, which together add up to a minimum level of background fluorescence $I_{bg}(x)$. We split this background fluorescence in two contributions: a contribution $I_{bg,0}$ that is constant over the entire image, originating from e.g. excitation light and ambient light, and a varying contribution caused by out-of-focus fluorescence, $I_{bg,z}$. Considering the membrane signal $I(x)$ of a particular GUV, the most dominant contribution to $I_{bg,z}$ comes from the out-of-focus membrane fluorescence of that GUV, which is high in the GUV interior and low outside the vesicle. We therefore write $I_{bg,z}$ as a Heavyside step function, with the interior out-of-focus fluorescence depending on the z-resolution of the system $c$ (between 0 and 1, where 1 means good z-resolution, e.g. scanning confocal microscopy) as well as the membrane signal as:

$$I_{bg,z}(x) = \begin{cases} (1-c)A, & \text{if } x < r \\ 0, & \text{otherwise} \end{cases} \tag{3}$$

Summing up, we obtain the total noise contribution:

$$I_{noise}(x) = I_{cam}(x) + I_{bg,0} + I_{bg,z}(x) \tag{4}$$

Including the noise in the experimentally obtained signal, we write:

$$I(x) = Ae^{-\frac{(x-r)^2}{2\sigma^2}} + I_{cam}(x) + I_{bg,0} + I_{bg,z}(x) \tag{5}$$

From the fluorescent signal, various metrics can be calculated to report a membrane intensity. The membrane is first segmented by an inner radius $r_{in}$ and an outer radius $r_{out}$. For a one-dimensional profile, the software can then obtain the sum of the signal $\sum(I(x))$ between $r_{in}$ and $r_{out}$, given by:

$$\sum I(x) = \sum_{x=r_{in}}^{r_{out}} (Ae^{-\frac{(x-r)^2}{2\sigma^2}} + +I_{cam}(x) + I_{bg,0} + I_{bg,z}(x))\Delta x \tag{6}$$

where $\Delta x$ is the integration unit, typically the size of a pixel. Integration is done over a total number of $N$ pixels, defined by $N\Delta x = r_{out} - r_{in}$. Using $N$, we can simplify eq. (6) to:

$$\sum I(x) = NI_{bg,0} + \sum_{x=r_{in}}^{r_{out}} (Ae^{-\frac{(x-r)^2}{2\sigma^2}} + I_{cam(x)} + I_{bg,z}(x)))\Delta x \tag{7}$$

The average signal $\overline{I}(x)$ can be calculated from eq. (7) by dividing over the number of pixels:

$$\overline{I}(x) = I_{bg,0} + \frac{1}{N} \sum_{x=r_{in}}^{r_{out}} (Ae^{-\frac{(x-r)^2}{2\sigma^2}} + I_{cam(x)} + I_{bg,z})\Delta x \tag{8}$$

Since $I_{cam}$ is random, when integration is done over a large number of pixels, or large $N$, $\sum_{x=0}^{N} I_{noise}(x) \to 0$ leaving:

$$\sum I(x) = NI_{bg} + \sum_{x=r_{in}}^{r_{out}} (Ae^{-\frac{(x-r)^2}{2\sigma^2}} + I_{bg,z})\Delta x \tag{9}$$

$$\overline{I}(x) = I_{bg} + \frac{1}{N} \sum_{x=r_{in}}^{r_{out}} (Ae^{-\frac{(x-r)^2}{2\sigma^2}} + I_{bg,z})\Delta x \tag{10}$$

As can be seen from eq. (9) and eq. (10), both the summed membrane intensity and average membrane intensity are influenced by the background signal $I_{bg}$ and thus require background subtraction. In addition, we see the average signal is dependent on the number of pixels. This means that care should be taken when the number of pixels differs between vesicles, e.g. because vesicles have different sizes. Furthermore, it is important to note that when $N$ is small, e.g. using a narrow membrane segmentation, $I_{cam}$ can have a dominant contribution.

Another descriptor that is often used to quantify membrane fluorescence is the maximum intensity. While the maximum intensity is a parameter that is easy to extract, its value is affected by pixelation and noise effects. For a continuous signal without noise, the maximum is simply given by the signal amplitude $A$. However, because of pixelation in the image, we do not find the fluorescence intensity exactly at the membrane position, but at a position that is at maximum $x_p$ separated from $r$, where $x_p$ is the size of a pixel. For a signal without noise, the maximum intensity that is seen is given by:

$$I_{max} = Ae^{-\frac{x_p^2}{2\sigma^2}} \tag{11}$$

Assume we have a 100x objective with a pixel size of $60nm$ and a PSF of $240nm$, and assume that $\sigma$ is half the width of the PSF, then $I_{max} = 0.9A$. However, for a typical 10x objective with a pixel size of $600nm$ and a PSF of $1.2\mu m$, $I_{max} = 0.6A$. Resolution can thus have a serious impact on the maximum intensity determined from a one-dimensional profile. Moreover, random noise in the image can affect the measured membrane position. Suppose that we have 20% noise (with respect to $A$), we can calculate the possible shift in membrane position $\Delta x$ due to addition of this noise. The signal with noise is given by:

$$I(\Delta x) = Ae^{-\frac{\Delta x^2}{2\sigma^2}} + 0.2A \tag{12}$$

We then find the shift in membrane position by identifying the $\Delta x$ for which $I(\Delta x) = A$, finally yielding $\Delta x = 0.6\sigma$. For a noise level of 50%, this yields $\Delta x = 1.2\sigma$. Assuming a $\sigma$ of 2 pixels, noise can cause a shift of one or two pixels dependent on the imaging settings.

#### Two-dimensional intensity profile

When considering the two-dimensional intensity profile, two other effects of pixelation must be taken into account.

First, angular slices must have a minimal thickness to extract pixel intensities from the slice (fig. S4B). While thinner slices approach one-dimensional intensity profiles, the slice thickness decreases when getting closer to the vesicle center. We compute the minimal angular separation $\theta$ where the slice thickness is larger than a pixel at a radial distance $x$. With the circle perimeter at a radial distance $x$ given by $L = 2\pi x$, and the number of slices given by $N = 360/\theta$, the perimeter of an angular slice is $\Delta L = \dfrac{2\pi x}{N}$. For $\Delta L > 1$, we need $\theta > \frac{180}{\pi x}$. For a radial distance of 10 pixels, that is an angular separation of at least 6°. This effect is especially important for smaller vesicles or lower magnification images.

Second, the angular slice is drawn with four continuous lines, meaning that the edges of the mask run right through pixels(fig. S4C). This means that pixels that are partly outside the mask are weighed disproportionally into the average, while pixels partly inside the mask might be ignored. This effect is stronger for smaller slices. To get an idea of the number of pixels within the mask versus the number of pixels under the edge of the mask, we calculate the area and the perimeter of the angular mask. The area of the mask can be calculated by:

$$A_{slice} = \frac{\theta}{360}(A_{c,out} - A_{c,in}) \tag{13}$$

$$A_{slice} = \pi\frac{\theta}{360}(r_{out}^2 - r_{in}^2) \tag{14}$$

The perimeter, or the number of pixels at the borders of a slice can roughly be calculated by summing up the length of the four sides:

$$N_{edge} = 2(r_{out} - r_{in}) + \pi\frac{\theta}{360}(r_{out} + r_{in}) \tag{15}$$

Suppose we have a vesicle of $10\mu m$ radius. When imaged with an 100x objective with pixel size $60nm$ and a PSF of $240nm$, the membrane has an apparent thickness of 4 pixels, while the distance to the center of the vesicle is 150 pixels. Using an angular separation of 5 °and a total width of 12 pixels around the membrane, $r_{in} = 140$, $r_{out} = 160$, we obtain $A_{slice}/N_{edge} \sim 4$. This means that pixels at the edges have a relative small contribution, and thus that data can safely be extracted by integration. Problems arise when the size of the object in the image decreases. If we image the same vesicle with a 10x objective with pixel size $600nm$ and PSF $1.2\mu m$, we obtain a membrane width of 2 pixels and a radial distance of 15 pixels. Now, using a ring of 3 times the membrane width gives us $r_{in} = 12$ and $r_{out} = 18$, resulting in $A_{slice}/N_{edge} \sim 0.5$. Since edge pixels would weigh disproportionally large in this situation, it would be better to extract membrane intensities with one-dimensional extraction methods.
