## Supporting Figures for "DisGUVery: a versatile open-source software for high-throughput image analysis of Giant Unilamellar Vesicles"

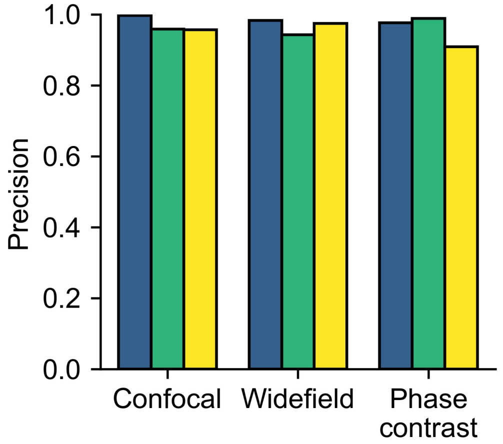


Figure S1: Precision of vesicle detection for different imaging types calculated from the same performance analysis results as shown in fig. 2G.

Table S1: Population sizes of vesicles in the different subcategories as counted by the four different observers in fig. 2H. The last column indicates the population size averaged over all observers.

| Category | Observer 1 | Observer 2 | Observer 3 | Observer 4 | average |
|---|---|---|---|---|---|
| Standard | 93 | 84 | 103 | 89 | 92 |
| Edge | 59 | 59 | 63 | 68 | 62 |
| Unsharp | 17 | 13 | 14 | 18 | 16 |
| Anomalous | 15 | 61 | 13 | 32 | 30 |

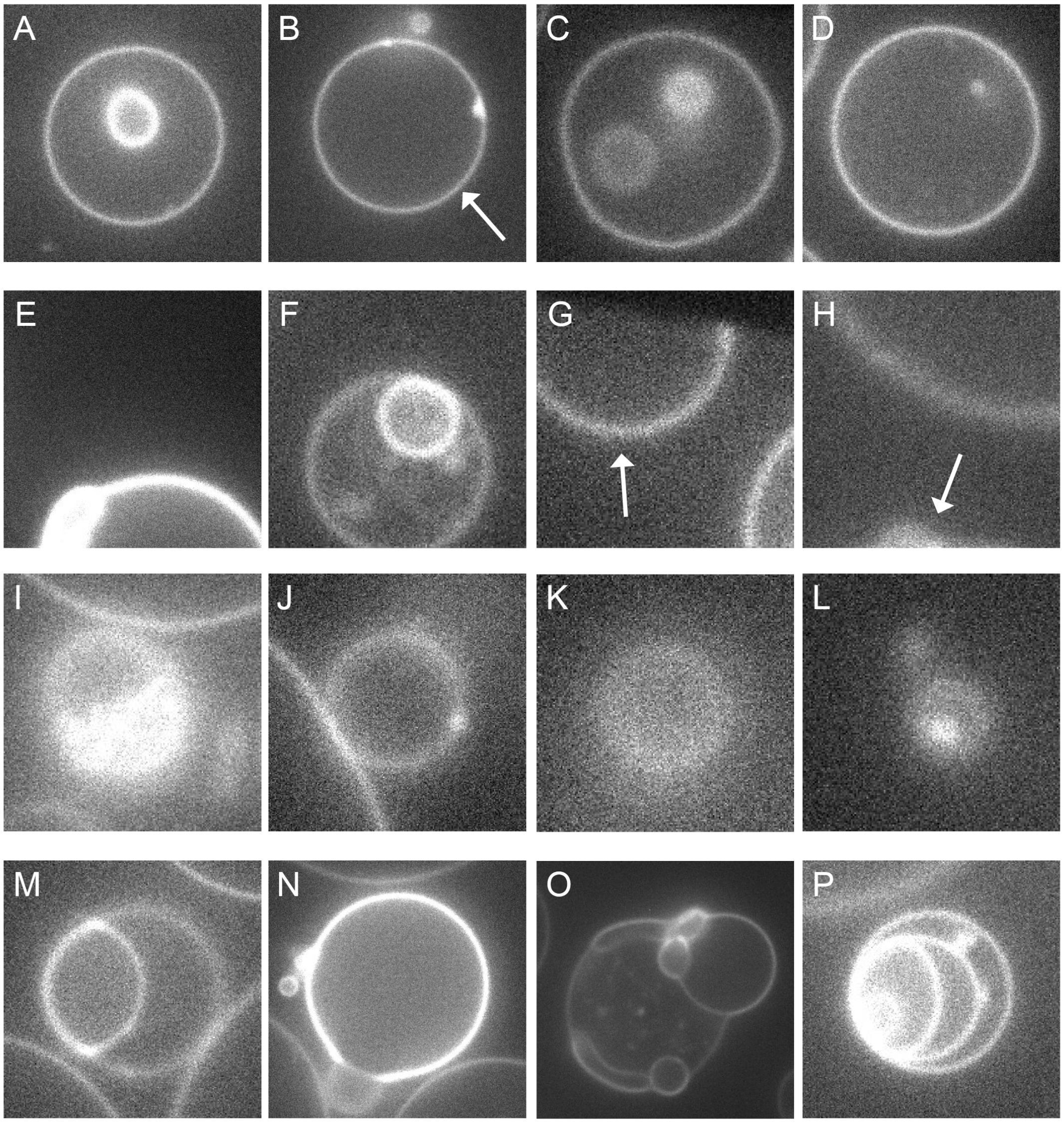


Figure S2: Gallery of example vesicles from different subcategories. In images containing multiple vesicles, the example vesicle has been indicated with an arrow. (A-D) Standard vesicles. (E-H) Vesicles at the edge of the image. (I-L) Vesicles that are out of focus. (M-P) Anomalous vesicles.

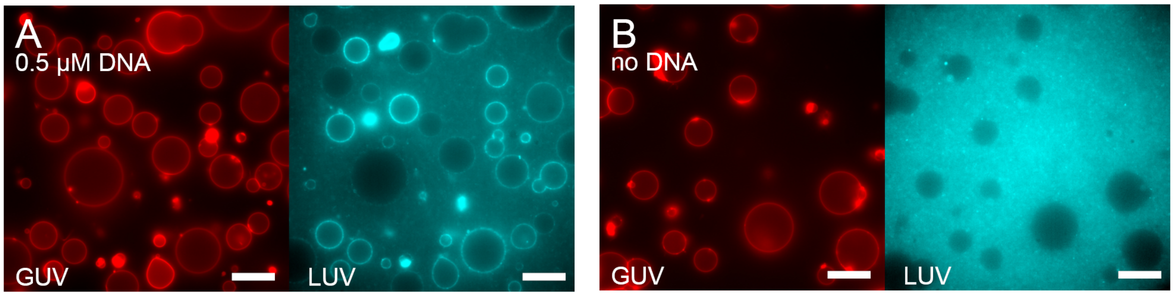


Figure S3: Binding of LUVs at different DNA concentrations. Images are epifluorescence images of Atto488 DOPE incorporated in the GUV membrane (red) and Atto 655 DOPE in the LUV membrane (cyan). Scale bar is 20 $\mu m$ in all images. (A) At $0.5\mu M$ cholesterol-DNA, LUVs bound to the GUV membrane. (B) No membrane localization was observed in absence of cholesterol-DNA.

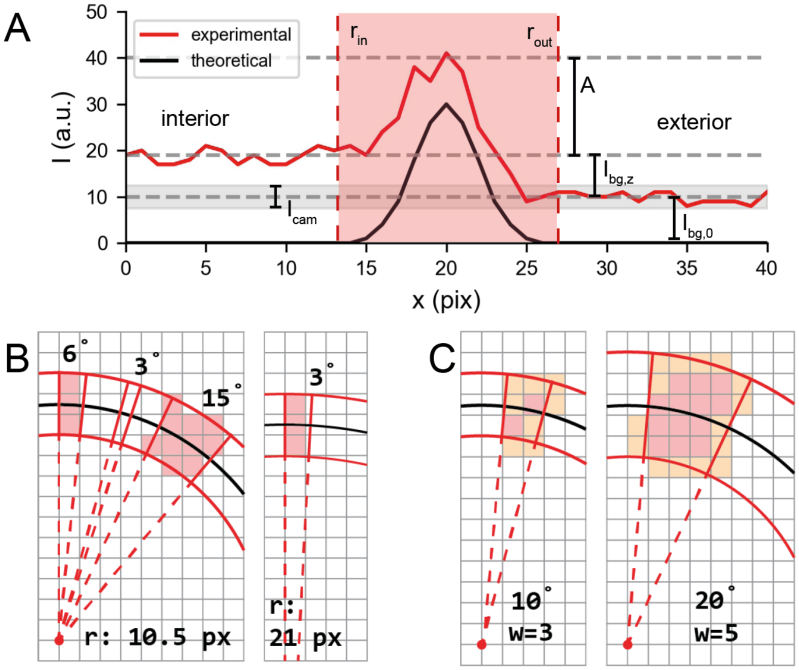


Figure S4: Experimental artefacts in membrane analysis. (A) Theoretical (black) and predicted experimental (red) one-dimensional membrane intensity profile of a GUV. The experimental profile is calculated with eq. (5) and eq. (3). While the theoretical profile is a simple Gaussian, the experimental profile is subjected to noise of various origins. Here, the radius $r = 20px$, $\sigma = 2px$, $A = 30$, $I_{cam} = 5$, confocality factor $c = 0.7$, and $I_{bg,0} = 10$. The red shaded part denotes the segmented area based on $r_{in} = 13$ and $r_{out} = 27$. (B,C) Discretisation effects in two-dimensional signal analysis. Segmentation lines are shown in red, the GUV membrane in black. (B) The apparent GUV size in the image imposes a minimum angular separation for integration. Integration should not be done with slices smaller than the pixel size. For a small GUV with a radius of 10.5 pixels, the minimum $\theta$ is 6°(left), while a more precise angular separation of 3° can be used for a vesicle twice as large (right). (C) Edge effects impact intensity analysis. Pixels located at the edge of the segmentation area (yellow pixels) are more prominent for smaller segmentation areas (left) as compared to larger regions (right). The size of the segmentation area is defined by the width of the ring as well as the angular separation.
